# Astrocyte TrkB.T1 deficiency disrupts glutamatergic synaptogenesis and astrocyte-synapse interactions

**DOI:** 10.1101/2024.10.22.619696

**Authors:** Beatriz T.C. Pinkston, Jack L. Browning, Michelle L. Olsen

## Abstract

Perisynaptic astrocyte processes (PAPs) contact pre- and post-synaptic elements to provide structural and functional support to synapses. Accumulating research demonstrates that the cradling of synapses by PAPs is critical for synapse formation, stabilization, and plasticity. The specific signaling pathways that govern these astrocyte-synapse interactions, however, remain to be elucidated. Herein, we demonstrate a role for the astrocyte TrkB.T1 receptor, a truncated isoform of the canonical receptor for brain derived neurotrophic factor (BDNF), in modulating astrocyte-synapse interactions and excitatory synapse development. Neuron-astrocyte co-culture studies revealed that loss of astrocyte TrkB.T1 disrupts the formation of PAPs. To elucidate the role of TrkB.T1 in synapse development, we conditionally deleted TrkB.T1 in astrocytes in mice. Synaptosome preparations were employed to probe for TrkB.T1 localization at the PAP, and confocal three-dimensional microscopy revealed a significant reduction in synapse density and astrocyte-synapse interactions across development in the absence of astrocytic TrkB.T1. These findings suggest that BDNF/TrkB.T1 signaling in astrocytes is critical for normal excitatory synapse formation in the cortex and that astrocyte TrkB.T1 serves a requisite role in astrocyte synapse interactions. Overall, this work provides new insights into the molecular mechanisms of astrocyte-mediated synaptogenesis and may have implications for understanding neurodevelopmental disorders and developing potential therapeutic targets.

## INTRODUCTION

Brain-derived neurotrophic factor (BDNF) and its receptor tropomyosin receptor kinase B (TrkB) constitute a powerful molecular partnership that serve to govern neuronal survival and maturation as well as synapse formation and plasticity; thereby shaping the intricate neuronal circuits that underlie mammalian behavior. While the role of BDNF/TrkB signaling in the development and maintenance of synapses has been extensively studied, we recently unveiled an unexpected player in this molecular pathway – the astrocyte.

We have demonstrated that astrocytes predominantly and nearly exclusively express a truncated isoform of the TrkB receptor, TrkB.T1 [1]. Further, evaluation of published sequencing data indicates that TrkB.T1 is the primary isoform identified in the cortex of both mice[2, 3] and humans[4, 5], and astrocytes across species are the highest expressers of TrkB[6] suggesting that astrocytes are a major recipient of BDNF in cortex. Astrocytes are a major glial cell type that have emerged as critical players in synapse development and stabilization. These functions are facilitated by their morphological complexity, which is best described as a dynamic spongiform shape, made up by thousands of fine, leaflet-like processes, often referred to as perisynaptic astrocyte processes (PAPs) that infiltrate the neuropil and allow a singular mammalian astrocyte to contact up to 100,000 synapses at a time[7]. From a functional standpoint, PAPs isolate synapses by forming a physical barrier that facilitates the maintenance of both ion [8–10] and neurotransmitter [11] in the synaptic space. PAPs are enriched with numerous transporters, receptors, and ion channels, such as the glutamate transporters (GLT1) and (GLAST) [12], the GABA transporter (GAT3) [13], and the inwardly rectifying potassium channel-4.1 (Kir4.1) [10, 14]. Together these proteins work to remove neuronally derived neurotransmitter and K^+^ ions from the synaptic cleft following neuronal activity [8, 15]. Astrocytes are also key players in the development and maturation of synapses through their expression of critical cell adhesion molecules (CAMs) and secretion of synaptogeneic cues [16]. Astrocytes have been increasingly implicated in the control of behavior[17–19], yet the underlying molecular mechanisms that guide astrocytic processes to synapses, and bidirectional neuron-astrocyte signaling that mediates astrocyte-synapse interactions and their consequent effects on behavior, remain to be fully elucidated.

Herein, utilizing a reduced astrocyte-neuron co-culture model system, our data indicate TrkB.T1 deficient astrocytes fail to form PAPs at synapses, suggesting that activation of the astrocyte TrkB.T1 receptor contributes to glutamatergic synapse recognition and subsequent PAP involvement. Analysis of cortical synaptosomes, which effectively capture both synapses and their surrounding perisynaptic astrocyte processes (PAPs), reveals a significant concentration of TrkB.T1. This enrichment is notably absent in mice lacking TrkB.T1, providing strong evidence for the localization of TrkB.T1 within the PAP. Furthermore, using a combination of protein and mRNA approaches as well as three-dimensional confocal microscopy, we demonstrate that loss of TrkB.T1 in astrocytes results in reduced glutamatergic synapse formation and astrocyte-synapse interactions in the motor and barrel cortex. Altogether, our findings reveal astrocytes serve as a primary recipient of neuronally derived BDNF and provide novel insights into the role of astrocyte TrkB.T1 in modulating excitatory synapse formation, with important implications in neuronal development and plasticity. These insights underscore the necessity of considering the interplay between BDNF signaling, astrocyte function, and synaptic integrity, which may hold the key to understanding and addressing a range of neurodevelopmental and neuropsychiatric disorders.

## METHODS

### Animals

All experiments were performed according to NIH guidelines and with the approval by the Institutional Animal Care and Use Committee at Virginia Polytechnic Institute and State University. All animals were maintained on a reverse 12-hour light/dark cycle (lights on at 10 PM, lights off at 10 am). Food and water were available *ad libitum.* Wild-type *TrkB.T1^+/+^* and Knockout *TrkB.T1^-/-^*[20] C57/B6 mice were used for these experiments. These global TrkB.T1 mutant mice were generated by crossing a heterozygous *TrkB.T1^+/-^*male with a heterozygous *TrkB.T1^+/-^* female. Global TrkB.T1 mutants are referred to as global wildtype (TrkB.T1^+/+^) or global knockout (TrkB.T1^-/-^) in the paper.

Astrocyte-specific conditional knockout mice were generated in house. Aldh1l1-cre/ERT2 (Jackson Labs #029655) mice were crossed to TrkB.T1^fl/fl^ mice. To obtain experimental animals, homozygous TrkB.T1^fl/fl^ Cre^+^ males were crossed to homozygous TrkB.T1^fl/fl^ Cre^-^ females, or vice versa, a homozygous TrkB.T1^fl/fl^ cre^+^ female was crossed to a homozygous TrkB.T1^fl/fl^ cre^-^ male. This cross resulted in offspring that were all homozygous TrkB.T1^fl/fl^ and either Cre^+^ or Cre^-^ mice. TrkB.T1^fl/fl^ Cre^+^ and Cre^-^ mice were injected once daily with 4-OH tamoxifen between postnatal day (PND) 8-11, creating astrocyte specific Trkb.T1 conditional knock out mice and littermate, age matched control mice. Tamoxifen was made to stock concentration of 20 mg/ml in 1:19 ethanol to canola oil via sonication for 60 minutes at 50°C. Injections were administered intraperitoneally at 20 µL prior to knowing the genotype of each animal. Astrocyte-specific conditional TrkB.T1 transgenic mice are referred to as conditional wildtype (Aldh1l1-TrkB.T1 cWT) and conditional knockout (Aldh1l1-TrkB.T1 cKO) in the paper. *TrkB.T1^-/-^* and TrkB.T1^fl/fl^ mice were a generous gift from Dr. Lino Tessarollo.

### Primary Neuron-Astrocyte Co-Cultures

Neurons were cultured from global wildtype (TrkB.T1^+/+^) PND 0-1 pups according to [1, 21]. Briefly, following cortical dissociation, tissue was minced and enzymatically dissociated with a Papain Dissociation Kit (Worthington) following manufacturer’s instructions. After obtaining a cell suspension of 150 µl, microglia and oligodendrocytes were first removed with 10 µL incubation with Cd11b^+^ (Miltenyi Biotec) and Mbp^+^ (Miltenyi Biotec) microbeads for 10 min at 4°C. The flow through was collected. Utilizing Miltenyi Biotec’s Neuron Isolation Kit, neuronal populations were isolated by incubating the cell suspension with 15 µl of biotinylated antibodies for 10 min at 4°C followed by a 10 min incubation with 15 µl anti-biotin microbeads. The cell suspension was then applied to a prepped LS column and neurons were collected in the flow through of two washes. Neuronal cell number was determined and 1.0-1.25×10^5^ cells were plated on 1 mm glass coverslips (Electron Microscopy Sciences; Cat. #: 72290-03) in a 24-well plate (Falcon; Cat. #: 353047). The coverslips were poly-l-lysine treated and laminin-coated prior to plating. Neurons were maintained in neuronal maintenance media (Neurobasal media, 2 mM 1-Glutamax, and 1X B27). On the second day post-plating, 2 µM cytosine arabinoside (AraC; Sigma; Cat. #: C6645) was added to reduce non-neuronal contamination. On the third day post-plating, a media change was performed to remove AraC.

At DIV 3, astrocytes from PND 3-5 global wildtype (TrkB.T1^+/+^) or global knockout (TrkB.T1^-/-^) littermates were isolated and plated on top of the neurons at a density of 0.75-1.0 x 10^5^. Briefly, following cortical dissociation and after obtaining a cell suspension as described above, microglia and oligodendrocytes were first removed with 10 µl incubation with Cd11b^+^ and Mbp^+^ microbeads for 10 min at 4°C. The flow through was collected and astrocytes were then acutely isolated using anti-ACSA-2^+^ (astrocyte cell surface antigen 2) Microbead Kit (Miltenyi Biotec). The flow through cell suspension was incubated in 15 µl of FcR blocking buffer for 10 min at 4°C followed by 15 µl of ACSA-2 microbeads for 10 min at 4°C. The cell suspension was then applied to a prepped LS column and astrocytes were eluted from the column after two washes. Astrocyte cell number was determined, and astrocytes were then subsequently plated or added to the WT neuronal plates. Subsequent half media changes occurred every 3-4 days.

### Co-Culture Experiments

#### Time Points

Primary co-cultures were fixed at DIV 3 and 10. Cells were washed with cold PBS followed by fixation with cold 4% paraformaldehyde (PFA) for 15 minutes at room temperature. Cells were washed with cold PBS 3 times before storing or initiating immunofluorescence.

#### KCl

Primary co-cultures were treated with exogenous KCl at DIV 8-9. KCl was applied in warmed media to a final concentration of 10 mM for 24 hours. Equal volumes of warmed media were added to control cultures. Following KCl treatment, cells were washed with cold PBS followed by fixation with cold 4% paraformaldehyde (PFA) for 15 minutes at room temperature. Cells were washed with cold PBS 3 times before storing or initiating immunofluorescence.

### Primary Co-Culture Immunofluorescence

Co-cultures were fixed at DIV 3 and 10 or DIV 10-11 for KCl experiments. Following fixation, cells were incubated for 1 hr in Fish Gel Blocking Buffer (2% Fish Gel [Sigma-Aldrich G741-100G], 0.2% Triton-X [Sigma-Aldrich T9284] in PBS) at room temperature. Coverslips were then incubated with primary antibodies in diluted blocking buffer (1:3 Blocking Buffer in PBS) overnight at 4°C. Cells were then washed for 5 min (5 times) in cold PBS before being incubated with AlexaFluor 488, 546, and 647 secondary antibodies. Secondary antibodies were made in 1X PBS and cells were incubated in secondary antibodies for 1.5 hr at room temperature and in the dark. Cells were washed for 5 min (5 times) in cold PBS before being incubated in DAPI (1:1000) for five minutes before final wash in PBS. Coverslips were mounted with ProLong Gold antifade reagent (Life Technologies; Cat. #: P36930) on glass slides (Fisher Scientific; Cat. #: 12-550-15).

Astrocyte morphology was visualized with the membrane-associated protein Glast, and pre- and post-synaptic elements were visualized with VGluT1 and Homer, respectively. Prior to image acquisition, the experimenter was blinded to experimental conditions. Fluorescent images were acquired with a Nikon A1 confocal microscope. Primary antibodies used: rabbit Glast (Abcam; Cat. #: ab416, 1:5000), mouse Homer (Synaptic Systems; Cat. #: 160-011; 1:500), and guinea pig VGluT1 (Millipore; Cat. #: AB5905, 1:1000). Secondary antibodies used: AlexaFluor 488 (Invitrogen; Cat. #: A11008; 1:500), AlexaFluor 546 (Invitrogen; Cat. #: A10036; 1:500), and AlexaFluor 647 (Invitrogen; Cat. #: A21450, 1:500).

### *In vitro* perisynaptic astrocyte process and synapse evaluation

A machine learning algorithm was used on Imaris x64 9.8.2. for evaluation of PAPs and synaptic associations. The Imaris *Surface Module tool* was employed to create a 3D and segmented rendition of Glast+ astrocytes. The smoothing factor, indicated in step two of the module, was applied during the surface creation. In the same module, a machine learning algorithm was used to classify each segment on the astrocyte. The astrocyte was segmented based on the average diameter of randomly selected PAPs, which were defined as astrocyte elements that were high in Glast intensity relative to the rest of the membrane, round elements that colocalized with pre-or post-synaptic elements, or both. Due to the varying intensities and shapes that these elements form, blinded experimenters provided Imaris with numerous examples of PAP elements and then allowed the module to classify the entire astrocyte surface based on the parameters and examples fed into the program. This allowed for an unbiased classification of astrocyte segments. Once the astrocyte was segmented and reconstructed, a 3D distance-based approach was used to quantify PAP and synapse interactions. The *Spots Module* in Imaris was used to create spots for each synaptic element based on the average diameter of each synaptic puncta. Next, a distance-based approach was used to quantify pre- and post-synaptic elements within .500 µm of each other. This provided a quantification of co-localized synapses. Next, the co-localized puncta were isolated in Imaris and using a distance-based approach classification within the *Spots Module*, co-localized spots were classified into those that were within .750 µm of a PAP structure, and those that were not. Finally, we quantified the percentage of PAPs that contained a co-localized spots.

### *In vivo* astrocyte morphology and synapse immunofluorescence

Astrocytes in conditional wildtype (Aldh1l1-TrkB.T1 cWT) and conditional knockout (Aldh1l1-TrkB.T1 cKO) were fluorescently labeled via AAV.Lck-GFP virus – pAAV.GfaABC1D.PI.Lck-GFP.SV40 (Addgene Viral Prep # 105598-AAV5). Postnatal day 0-1 pups were intraventricularly injected with 2-3 uL 2.3 x 108 virus following hypothermia-induced anesthesia. The injection site was determined following Kim et al., 2014 [22] and as described previously by our lab [1] with equidistance between the bregma and lambda sutures, 1 mm lateral from the midline, and 3 mm depth. A 10 µL Hamilton syringe and 32G needle was used. Animals were left with their mother and subsequently injected with Tamoxifen once a day between PND 8-12 as described above. At time of collection, animals were anesthetized with peritoneal injections of 100 mg/kg of ketamine/xylazine and intracardially perfused with PBS followed by 4% PFA for 10 minutes. Brains were post-fixed overnight in 4% PFA before being transferred to a sucrose solution (30% Sucrose in PBS). Brains were stored at 4°C in sucrose until they sunk to the bottom of their conical tube (1-2 days). After sinking, brains were removed from sucrose, washed with cold PBS, and immersed in isopentane (Sigma; 2-Methylbutane; Cat. #: 320404) for 5-10 seconds. A small metal ramekin was half-filled with isopentane and placed on dry ice, allowing the isopentane solution to cool down to −45°C to enable tissue snap-freezing. Brains were then stored at −80°C for later sectioning or sectioned immediately on a Leica Micrtotome (Leica SM2010 R Sliding Microtome). Sections were 100 µm and stored in anti-freeze solution (30% glycerol, 30% ethylene glycol in 1X PBS) at −20°C.

### Immunofluorescence

100 µm coronal sections containing motor and somatosensory whisker barrel cortex were selected as detailed in Franklin and Paxinos. Sections were washed for 3 minutes (5 times) with cold PBS before being blocked for 1 hr in fish Gel Blocking Buffer (2% Fish Gel [Sigma-Aldrich G741-100G], 0.2% Triton-X [Sigma-Aldrich T9284] in PBS) at room temperature. Slices were then incubated with primary antibodies made in diluted blocking buffer (1:3 Blocking Buffer in PBS) overnight at 4°C. Following primary incubation, slices were washed for 5 minutes (5 times) in cold PBS before being incubated with AlexaFluor 555/546 and 647 secondary antibodies. Secondary antibodies were made in 1X PBS at a 1:500 concentration. Sections were incubated in secondaries for 1.5 hours at room temperature before being washed for 5 min (5 times) in cold PBS, incubated in DAPI (1:1000) for five minutes before final wash in PBS. Sections were finally mounted with ProLong Gold antifade reagent (Life Technologies; Cat. #: P36930) on glass slides (Fisher Scientific; Cat. #: 12-550-15) and covered with cover glass (Cat #: 12-548-5E, Fisher Scientific).

Primary Antibodies Used: Mouse PSD 95 (Abcam; Cat. #: AB2723; 1:500), Guinea Pig VGluT1 (Millipore; Cat. #: AB5905; 1:1000) and Rabbit VGluT2 (Abcam; Cat. #: AB216463; 1:1000). Secondary Antibodies Used: Goat anti-Guinea Pig 555 (Invitrogen; Cat. #: A21435; 1:500), Goat anti-Rabbit 546 (Invitrogen; Cat. #: A11010; 1:500), and Donkey anti-Mouse 647 (Invitrogen; Cat. #: A31571; 1:500).

### In vivo synaptosome preparations

Synaptosome isolations were completed as previously described [23]. PND 22-28 Aldh1l1-TrkB.T1 cWT Aldh1l1-TrkB.T1 cKO mice were euthanized via CO-2 and their brains rapidly dissected and cortices dissected in ice cold bubbled 1x Krebs buffer (1.26M NaCl, 25mM KCl. 250mM NaHCO3, 12mM NaH2PO4, 12mM MgCl2, 25mM CaCl2; pH 7.2). Cortices were added to 2ml Dounce Glass homogenizers (Wheaton, Cat#: 357422) on ice with 1.5 mL freshly made homogenizing buffer (2.5ml 4X gradient buffer (containing 109.54g Sucrose, 606mg Tris, 5mL 0.2M EDTA, pH 7.4), 7.5mL ddH2O, and 50mM dithiothreitol) and homogenized with 10-15 strokes. The resulting supernatant was transferred to a 2mL centrifuge tube and centrifuged at 1000xrcf for 10 minutes at 4°C. 100µL of the supernatant was taken and diluted in 300µL of 1x Krebs buffer with phosphatase (Sigma Aldrich, Cat. #: P5726) and protease (Sigma-Aldrich, Cat. #: P8340) inhibitors, this was stored as the whole cortex fraction. The remaining supernatant was transferred to another 2mL centrifuge tube and spun at 16000xrcf for 20 minutes at 4°C. The supernatant was kept as the cytosolic fraction. The remaining pellet was resuspended in 700µL of homogenizing buffer using a pipette and laid on top of a pre-prepared percoll (Sigma-Aldrich, Cat. #: P1644-100mL) gradient of 3%, 10%, 15%, and 23% from top to bottom in gradient buffer + 50mM dithiothreitol in a 4mL ultracentrifuge tube (Beckmann Coulter, Cat#: 349622). Tubes were placed in a chilled Beckman TLA 100.3 rotor and then placed in a Optima Max-XP ultracentrifuge (Beckman Coulter, Cat#: 393315) and spun at 31,000xg for 6 minutes at 4°C. Following ultracentrifugation, four bands and one pellet were identified. The third and fourth band from the top were collected using a glass Pasteur pipette as the synaptosome fraction. The resulting synaptosomes were transferred to a fresh 2 ml centrifuge tube and washed twice with 1x Krebs buffer at 15000xrcf for 15 minutes at 4°C, or until a solid synaptosome pellet was visualized. Once the synaptosomes pelleted, the supernatant was discarded, and the pellet was resuspended in 100µl of Krebs buffer plus inhibitors. All fractions protein content was assessed through a Pierce BCA assay, diluted to 2µg/µl in Krebs buffer plus inhibitors if needed, and stored at −80° until used for western blotting.

### Western Blot Evaluation

Sample lysates were diluted to a concentration of 0.33µg/µL with krebs buffer and mixed with 3.75µL of 4X Laemmli sample buffer (BioRad, Cat #: 161-0747) containing 200mM dithiothreitol. Samples were incubated for 10 minutes at 65°C. Equal amounts of protein were loaded into each lane of a 4–20% gradient precast acrylamide SDS-PAGE gel (Bio-Rad, Cat #: 4561096). Gels were loaded with 2µL of Precision Plus ProteinTM Standard Dual Color ladder (BioRad, Cat #: 161-0374), followed by 15µL of Hek cell lysate for negative control, and 15µL of each sample. Proteins were separated at 200 V constant. Gels were transferred utilizing the Trans-Blot® TurboTM RTA nitrocellulose transfer kit (Biorad, Cat # 170-4270), at 1.0A constant for 30 minutes. Following transfer, membranes were blocked in blocking buffer (1:1 ratio of TBS (150mM NaCl, 2.6mM KCl, 24.7mM Tris) to Interecept® Blocking Buffer (Li-Cor, Cat #: 927-60001). Blots were incubated in primary antibody solution containing 1:1 ratio of TBST (TBS + 0.002% Tween 20) to Intercept® Blocking Buffer overnight. The membranes were then rinsed three times for five minutes in TBST and then incubated with fluorescently conjugated secondary antibodies for one hour. Blots were once again washed three times for five minutes and imaged under a LiCor Odyssey Scanner 9120 Classic. Images were later analyzed under Image Studio, normalizing samples signal mean to area first then to in lane housekeeping genes, and finally to the sham group. Primary antibodies used were Rabbit Homer (1:5000, Protein Tech, Cat#: 12433-1-AP), Rabbit Kir4.1(1:5000, Protein Tech, Cat. #: 12503-1-AP), Rabbit TrkB (1:5000, Millipore, Cat. #: 07-225), Rabbit Glt1 (1:5000, Millipore, Cat. #: AB1783), Rabbit GRIN2B (1:5000, Protein Tech 21920-1-AP), Guinea Pig Vglut1 (1:5000, Millipore, Cat. #: AB5905), Mouse MAP2 (1:2500, Sigma, Cat. #: M1406), and Mouse Actin (1:10000, Millipore, Cat. #: A5441) Secondaries utilized were Goat anti mouse 680 (1:10000, Li-Cor, Cat. #: 925-68070), Donkey anti Guinea Pig 680 (1:10000, Li-Cor, Cat. #: 926-68077), Donkey anti Mouse 800 (1:10000, Li-Cor, Cat. #: 926-32212) and Goat anti Rabbit 800 (1:10000, Li-Cor, Cat. #: 925-32211).

### RNA Isolation and qPCR

Samples from our three developmental timepoints of interest were collected from wildtype (Aldh1l1-TrkB.T1 cWT) and conditional knockout (Aldh1l1-TrkB.T1 cKO). One paraformaldehyde-fixed, 100 µm thick coronal section was used for each animal (See *“in vivo astrocyte morphology and synapse immunofluorescence”* above for details on brain slicing). Two 750 µm tissue punches (one per hemisphere) containing the somatosensory cortex were collected. RNA was extracted using the manufacturer (Life Technologies, Carlsbad, CA, USA)-recommended procedures for a RecoverALL Total Nucleic Acid Isolation Kit (Cat. #: AM1975) optimized for paraffin-embedded (FFPE) or paraformaldehyde exposed samples. Following RNA isolation, 200 ng of RNA was reversed transcribed into cDNA using BioRad’s iScript SuperMix (Cat. #: 1708841). All cDNA was normalized to 350 ng following conversion. The relative mRNA expression levels were determined using real-time quantitative PCR by Taqman Fast Advanced Master Mix for qPCR (Cat. #: 4444557) and TaqMan specific probes. The ddCt method was used to quantify relative mRNA expression levels, with each normalization indicated where appropriate.

### *In Vivo* Imaging and Analysis

#### Synaptic Puncta in Astrocyte Regions

For imaging astrocyte PAP and synapse histology, Z-stack confocal images were acquired from layer II/III motor cortex and layer IV whisker barrel cortex using a Nikon A1 confocal microscope with 60X/NA 1.4 oil immersion lens and at 5X digital zoom. eGFP-expressing astrocytes and the synaptic puncta within the astrocytes were obtained from both hemispheres. The astrocytes were sampled at a 2048×2048 frame size, 0.225 µM step size, and a 3 µM-total Z-Stack was imaged and generated. Two to three images from each hemisphere and both hemispheres were imaged per animal for a total of three to four images per animal. Imaris (Bitplane, Zurich, Switzerland x64 9.8.2) was used to quantify synaptic punctate at a three-dimensional resolution according to the published literature [24–28]. To examine synapses within an astrocytic region, eGFP+ astrocytes were located and a 10 µm x 10 µm x 3 µm (100 µm^3^) region of interest (ROI) at least 10 µm from the cell body and closer to the distal boundaries of the astrocyte territory was randomly selected. Using the *Imaris Surface module*, a surface reconstruction of the eGFP+ astrocyte leaflets was built. For Spot Detection, we followed similar procedures as described before [24–28]. Briefly, the pre-synaptic (VGluT1 and VGluT2) and postsynaptic (PSD95) punctate were detected using the *Imaris Spot Module* with 0.4 µm and 0.3 µm diameters respectively, based on both the published literature and by manually finding the average diameter of our punctate across various samples. Only synaptic puncta/spots inside the astrocyte-object were quantified. We determined how many pre-synaptic spots were directly opposed to postsynaptic spots with the distance of 0.750 µm. The spots module and distance-based classifications enabled Imaris to quantify the total number of pre-post- and colocalized puncta within our ROIs.

#### Astrocyte Morphology and Astrocyte Synapse Interactions

Layer II/III motor cortex and layer IV whisker barrel cortex astrocytes were imaged on a Nikon A1 confocal with 60x oil immersion lens and 2X digital zoom. Laser power and gain were adjusted for each astrocyte and corresponding synaptic puncta. eGFP-expressing astrocytes that were visually isolated from neighboring astrocytes were sampled at a 2048×2048 frame size, 1 µm Z-step size, and 2x digital zoom. Care was taken to ensure that the entirety of each astrocyte in the Z-plane was imaged. After acquisition, Z-stacks were 3D reconstructed and analyzed on Imaris x64 V.V.V. The *Surface Module* was utilized to create a surface reconstruction and to estimate astrocyte volume. Astrocyte-synapse interaction analysis was estimated as described [29–32]. Briefly, the *Surface Module* in Imaris was used to generate a 3D reconstruction of each astrocyte, using the Lck-GFP channel signal to build the build the isolated surface. A special masked channel was generated using the 3D-surface rendering to isolate the Lck-GFP expressing astrocyte from Lck-GFP background and other astrocytes. The masked channel was then utilized to perform colocalization analysis within Imaris. Colocalization between the masked Lck-GFP and the Alexa 546 or Alexa 647 signal, representing VGluT1/VGluT1 and PSD95 markers respectively. To remove the background of the synaptic signals, repeated measurements of unambiguous puncta of fluorescent signal intensity across multiple optical slices were obtained. The average of these measurements was measured to threshold the synaptic channels. Next, the masked Lck-GFP channel was selected as a region of interest to focus on, and a new colocalization channel was generated which quantified the percentage of ROI (astrocyte eGFP signal) colocalized with the synaptic channel. This allowed for assessing the co-registry of fluorescent astrocyte and synapse signals within a given voxel and served as a proxy of astrocyte/leaflet proximity to pre- and post-synaptic elements.

### Statistical Analysis

Statistical significance and figures were generated using GraphPad Prism 10.1.2. All data are represented as mean +/- SEM, with n’s indicated as appropriate. The Shapiro-Wilk Test was performed to determine the normality distribution of each data set, and outliers were determined using GraphPad Prism’s ROUT method with a Q = 5%. For data where only one comparison was needed, student’s t-tests were performed. Two-way ANOVAs followed by Turkey’s post-hoc tests were performed for all data with multiple comparisons.

## RESULTS

### Development of astrocyte-synapse interactions are mediated via TrkB.T1 receptor *in vitro*

Given the canonical role of BDNF and TrkB in synapse formation and plasticity, we sought to investigate the role of astrocyte TrkB.T1 in synapse formation *in vitro*. Previously, we demonstrated in an astrocyte-neuron co-culture model system that TrkB.T1^-/-^ KO astrocytes fail to support the formation of structural or functional glutamatergic synapses in WT neurons, indicating a potential role of TrkB.T1 in the formation of synapses [1]. To directly investigate the underlying mechanisms by which astrocyte TrkB.T1 regulates synapse formation and maintenance, and how disruption of this process may contribute to the synaptic deficits observed, we examined the formation of PAPs and their interactions with synapses using primary astrocyte-neuron co-cultures from TrkB.T1^+/+^ WT and TrkB.T1^-/-^ KO mice. Cortical TrkB.T1^+/+^WT neurons were cultured from P0-1 mouse pups and allowed 3 days to recover. Subsequently, TrkB.T1^+/+^ WT or TrkB.T1^-/-^ KO astrocytes were plated on top of the TrkB.T1^+/+^ WT neuron cultures. After maintaining co-cultures for 10 days in vitro (DIV), co-cultures were paraformaldehyde fixed and confocal images of excitatory synapses (VGluT1+/Homer+) and astrocytes (GLAST+) were acquired **(Figure 1A)**. Immunolabeling for the astrocyte specific glutamate aspartate transporter (GLAST), which served as a membranous label as opposed to a cytoskeletal label for astrocytes, revealed interactions between astrocytes and neurons, with astrocytes overlapping neuronal branches and extending finer ramifications into synaptic sites. We additionally observed astrocytes formed high-intensity GLAST+ hotspots that appeared to co-localize and/or sometimes appeared to cradle pre-synaptic (VGluT1) and post-synaptic (Homer) puncta **(Figure 1A)**. Notably, these structures were significantly reduced in cultures containing TrkB.T1^-/-^ KO relative TrkB.T1^+/+^ WT cultures **(Figure 1B)**. Given the decrease in neuronal synapses observed when neurons are cultured with TrkB.T1^-/-^ KO astrocytes [1], we aimed to examine the role of BDNF/TrkB.T1 astrocyte signaling in mediating synapse formation and astrocyte-synapse associations. To examine the temporal dynamics of synapse and PAP development *in vitro*, and the necessity of TrkB.T1, we paraformaldehyde fixed co-cultures at different timepoints. We employed a machine learning algorithm in Imaris to both identify and classify the *in vitro* PAPs, these were defined as spots characterized by high intensity GLAST+ labeling, made of a “donut” shape, or both. We henceforth refer to these PAPs as “Astrocyte Elements.” Imaris classifications then allowed for the quantification of astrocyte elements containing active synapses; active synapses were defined as spots of co-localized pre- and post-synaptic puncta, obtained using the Imaris *Spots Module* **(Figure 1C)**. We established that synapses peaked at DIV10 *in vitro*, and thus quantified the upregulation of synapses between DIV 3 and 10. Two-way ANOVA revealed a significant effect of genotype (F(1, 13) = 42.54, p<0.0001; N = 4 DIV 3 WT, N = 4 DIV 3 KO, N = 5 DIV 10 WT, N = 4 DIV 10 KO, with n = 3-5 astrocytes per culture; **Figure 1D**), as well as a significant interaction (F(1, 13) = 13.37, p = 0.0029) with development (F(1, 13) = 10.70), p = 0.0061). *Post hoc* Turkey’s multiple comparison tests indicated that the percentage of astrocyte elements engaged with an active synapse significantly increase between DIV 3 and 10 in TrkB.T1^+/+^ WT (p < 0.05) but not in TrkB.T1^-/-^ KO conditions (p = 0.1634). While there were no differences between TrkB.T1^+/+^ WT and TrkB.T1^-/-^ KO conditions at DIV 3 (p = 0.2462), at DIV 10, TrkB.T1^+/+^ WT co-cultures had significantly more astrocyte-synapse interactions relative to TrkB.T1^-/-^ KO co-cultures (p <0.0001) **(Figure 1D-E)**. As expected, the Imaris *Spot Module* tool analysis, which enables the quantification of synapses [26], revealed that the number of synapses also significantly increase between DIV 3 and 10 in TrkB.T1^+/+^ WT (p = 0.0003) but not in TrkB.T1^-/-^ KO conditions (p = 0.65). At DIV 10, there is a significant difference between TrkB.T1^+/+^ WT and TrkB.T1^-/-^ KO conditions (p = 0.0001; **Figure 1F-G**). Altogether, we found a significant effect of genotype (F(1, 13) = 27.98, p<0.0001; N = 4 DIV 3 WT, N = 4 DIV 3 KO, N = 5 DIV 10 WT, N = 4 DIV 10 KO, with n = 3-5 astrocytes per culture; **Figure 1F-G**) as well as a significant interaction (F(1, 13) = 12.84, p = 0.003) with development (F(1, 13) = 19.68, p = 0.0007; **Figure 1F-G**).

**Figure 1:**
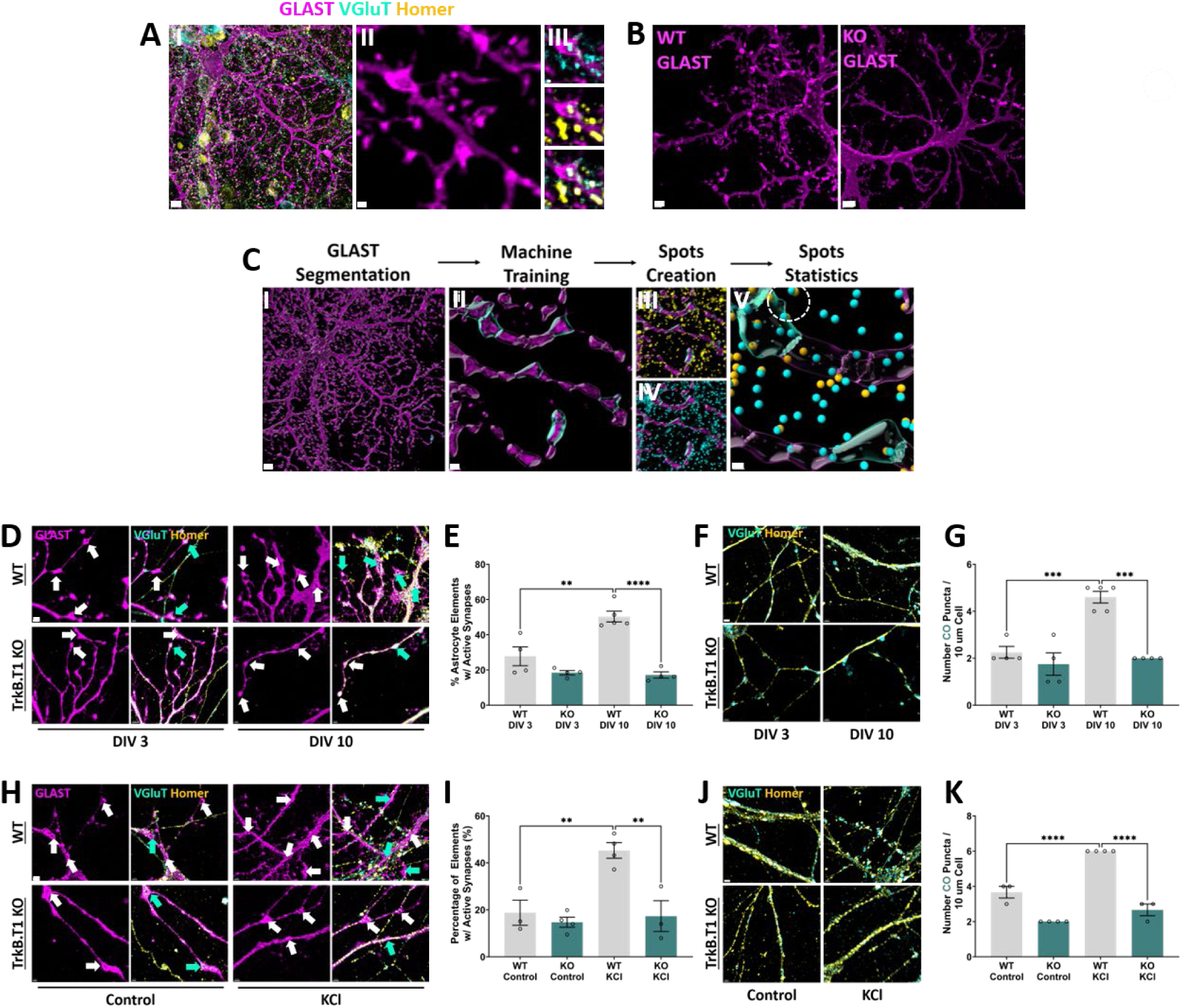
Astrocytes interact with developing neuronal synapses via TrkB.T1 *in vitro*. **(A)** *(I)* Representative image of GLAST+ astrocyte co-cultured with neurons. Scale Bar = 10 μm *(II)* Magnified astrocyte branches depicting high-intensity and/or cradle-like structures, termed PAP elements. Scale Bar = 1 μm. *(III)* These PAP elements were found to be associated with colocalized VGluT and Homer puncta. Scale Bar = .5 μm. **(B)** Representative images of GLAST+ TrkB.T1+/+ WT and TrkB.T1-/- KO astrocyte. TrkB.T1-/- KO astrocytes appeared to have a decrease in PAP elements. Scale Bar = 10 μm. **(C)** Representative images of Imaris workflow to quantify astrocyte-synapse interactions in vitro. (I-II) First, GLAST+ astrocyte is masked and segmented using Machine Learning classification to both identify and quantify PAP elements. Scale Bar = I: 15 μm; II: 2 μm (III-IV) Imaris Spots Creation enabled for the ability to both identify and quantify pre-(VGluT+) and post-synaptic (Homer) puncta. Scale Bar = 2 μm. (V) Analysis involved quantifying the percentage of PAP elements that contained an active synapse, defined as distance based co-localization of VGluT and Homer Puncta. White dashed circles indicate a PAP associated with active synapse. Scale Bar = 1 μm. **(D)** Representative images of TrkB.T1+/+ WT and TrkB.T1-/-KO astrocytes co-cultured with TrkB.T1+/+ WT neurons. Images depict the maturation of GLAST+ branches associated with neuronal processes between DIV 3 and DIV 10. White arrows indicate segments selected by Imaris as “Astrocyte Elements.” Teal arrows indicate Astrocyte Elements that contained a co-localized pre- and post-synaptic puncta, an “Active Synapse.” Scale Bar = 2 μm. **(E)** Quantification of the percentage of PAPs that contained an active synapse between DIV 3 and DIV 10. **(F)** Representative images of synapses from TrkB.T1+/+ WT neurons in the presence of WT astrocytes or TrkB.T1-/- KO astrocytes across development. Scale Bar = 2 μm. **(G)** Quantification of synapse numbers between DIV 3 and DIV 10. **(H)** Representative images of TrkB.T1+/+ WT and TrkB.T1-/- KO GLAST+ astrocyte branches associated with neuronal processes in control or 10 mM KCl-treated media. White arrows indicate segments selected by Imaris as “Astrocyte Elements.” Teal arrows indicate Astrocyte Elements that contained a co-localized pre- and post-synaptic puncta, an “Active Synapse.” Scale Bar = 2 μm. **(I)** Quantification of the percentage of PAP elements that contained an active synapse after KCl treatment relative no-treatment controls. **(J)** Representative images of synapses from TrkB.T1+/+ WT neurons in the presence of TrkB.T1+/+ WT or TrkB.T1-/- KO astrocytes in control or 10 mM KCl-treated media. Scale Bar = 2 μm. Data represented as averages ± SEM. Mixed model ordinary two-way ANOVA with Turkey’s multiple comparisons test. Each dot represents an individual culture with n = 3-5 cultures and n = 5 cells per culture (E and J) and15 ROIs per cell (G and K). *p < 0.05, **p < 0.001, *** p < 0.001.

### TrkB.T1 KO astrocytes fail to respond to increased neuronal activity *in vitro*

During early development, BDNF levels are high as neurons finalize their migration, survival, and engage in synaptogenesis [33]. Later in development, levels of BDNF, particularly in the cortex, are reported to decrease as its release becomes more specific, with high neuronal activity, learning and memory, and experience-dependent plasticity catalyzing more localized BDNF production[1, 34]. To test if TrkB.T1 is also involved in the development of synapses and astrocyte-synapse interactions in the context of activity-dependent release of BDNF, we treated co-cultures with 10 mM KCl for 24 hours. Elevated extracellular KCl (10 mM) has been shown to induce neuronal depolarization, concomitant neuronal activity [35] as well as increase BDNF expression in cultured neurons [36]. Next, we co-cultured neurons and astrocytes as above. Control media or media with high K+ (10 mM) was delivered at DIV 8-9 before paraformaldehyde fixation 24 hours after treatment. Imaris analysis and a mixed model effect analysis revealed a significant difference between TrkB.T1^+/+^ WT and TrkB.T1^-/-^ KO conditions (F(1, 10) = 14.25, p = 0.0036; N = 3 WT Control, N = 4 WT KCl, N = 4 KO Control, N = 3 KO KCl; **Figure 1H-I**), as well as a significant interaction (F(1, 10) = 7.978, p = 0.018) and effect on treatment (F(1, 10) = 11.81, p = 0.0064). Post hoc test results revealed that the percentage of astrocyte elements engaged with an active synapse significantly increase after KCl treatment in TrkB.T1^+/+^ WT (p = 0.0013) but not in TrkB.T1^-/-^ KO conditions (p = 0.6742). In KCl treatments, there is also a significant difference between TrkB.T1^+/+^ WT and TrkB.T1^-/-^ KO conditions, with TrkB.T1^+/+^ WT conditions having significantly higher astrocyte-neuron interactions (p = 0.0009) **(Figure 1H-I)**. Imaris Spots analysis similarly revealed a significant increase in synapses after KCl treatment in TrkB.T1^+/+^ WT (p < 0.0001) but not in TrkB.T1^-/-^ KO conditions (p = 0.1420) (**Figure 1J-K**). At both baseline and KCl treatments, paired multiple comparisons tests also revealed higher synapses in TrkB.T1^+/+^ WT relative to TrkB.T1^-/-^ KO conditions (p = 0.0007; p < 0.0001; **Figure 1J-K**). Together, our results using a simplified neuron-astrocyte model system demonstrate that astrocyte TrkB.T1 deficiency disrupts astrocyte-synapse interactions across *in vitro* development and that astrocyte TrkB.T1 serves a requisite role in astrocyte synapse interactions in the context of activity-dependent plasticity. These findings raise intriguing questions about the precise localization of TrkB.T1 within astrocytes, and whether it is potentially enriched in PAPs.

### TrkB.T1 is enriched at the synapse *in vivo*

To further elucidate the role of TrkB.T1 in astrocyte-synapse interactions and validate our *in vitro* observations, we next sought to investigate the subcellular localization of TrkB.T1 *in vivo*. Specifically, we aimed to determine whether TrkB.T1 is indeed concentrated at PAPs and synaptic interfaces in the brain, which would provide support for its proposed function in mediating astrocyte-synapse communication.

To achieve TrkB.T1 knock out in astrocytes, homozygous TrkB.T1^fl/fl^-Aldhl1Cre^+^ or TrkB.T1^fl/fl^-Aldhl1Cre^-^ were injected with 20 ul (0.2 mg) 4-OH tamoxifen once daily[1, 37] between PND8-11. This protocol results in near complete elimination of TrkB.T1 gene and protein, with small effects of TrkB.FL at the gene, but not the protein level (**Supplemental Figure 1**). To specifically address the localization of TrkB.T1 at the synapse, we employed a synaptosome preparation. Synaptosome preparations are versatile tools for studying synapses; they contain intact pre- and postsynaptic membranes along with attached PAPs, enabling structural, functional, and biochemical analyses of the synaptic machinery and its components [23]. Studies have demonstrated that these preparations collect PAP-associated proteins [38]. Drawing from our earlier RNA sequencing findings [21] and publicly available datasets[2–5], we established that TrkB.T1 exhibits high expression in mammalian cortex. Consequently, we anticipated that our synaptosome samples would be enriched in TrkB.T1. We validated our synaptosome preparations through electron microscopy and western blotting for synaptic and cytosolic proteins (**Figure 2A-B**). In the control WT samples, the postsynaptic protein Homer was enriched 4.43-fold more in the synaptosome (SYN) lysates than in the whole-brain (WB) lysates and 8.09-fold more strongly relative to the cytosol (CYT) lysates (**Figure 2B-C**). The protein expression level of neuronal microtubule-associated protein-2 (MAP2) was 8.87-fold greater in CYT lysates than in SYN lysates and 2.96-fold greater than in WBs (**Figure 2B and E**). In addition, we found high protein expression levels of astrocyte GLT1 (**Figure 2D**), TrkB.FL (**Figure 2F**), and TrkB.T1 (**Figure 2G**), indicating that the synaptosome preparations effectively pulled down the tri-partite synapse and its associated proteins, including TrkB.

**Figure 2:**
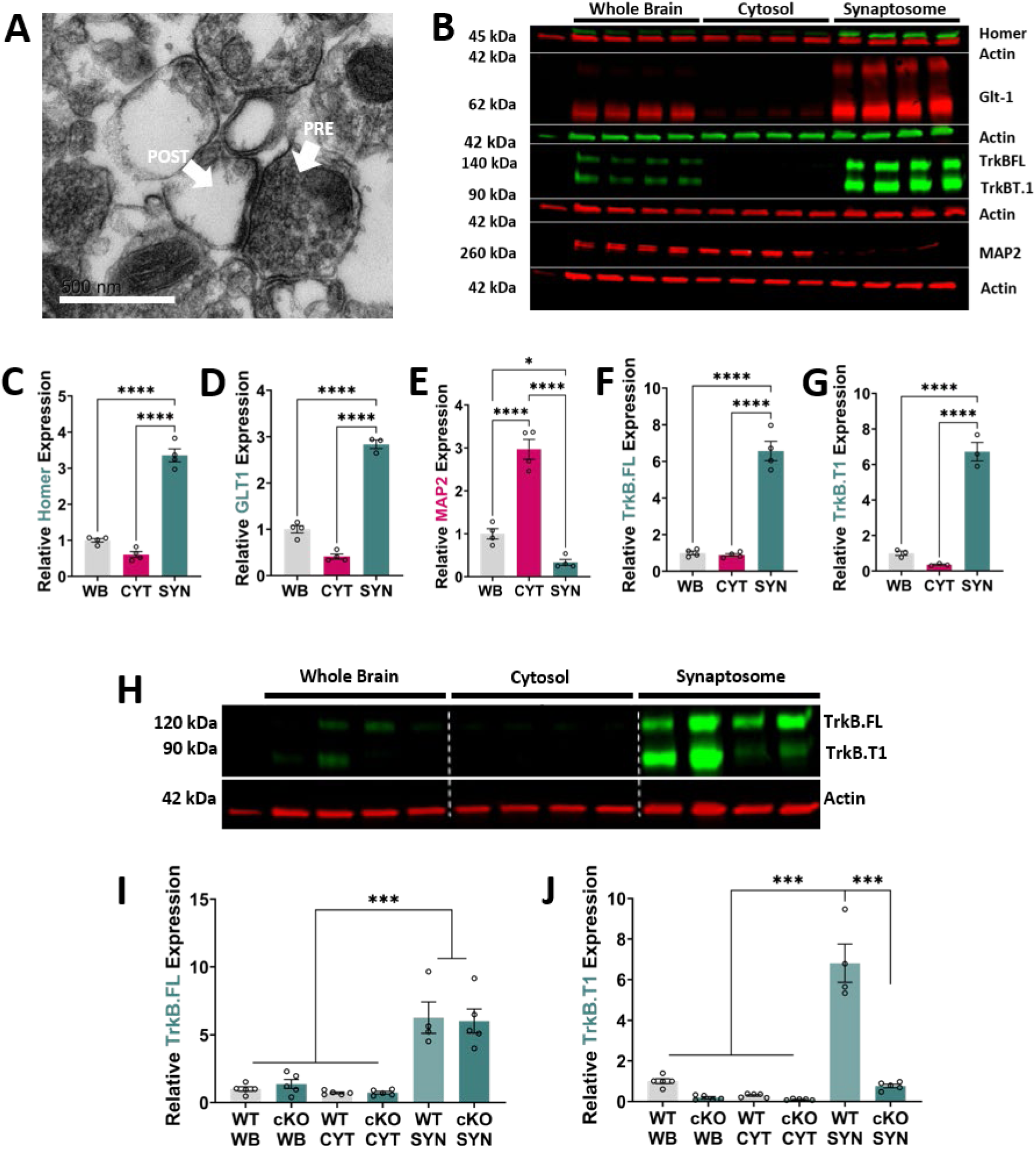
TrkB.T1 protein is enriched in perisynaptic astrocyte processes *in vivo*. **(A)** Representative image of transmitted electron micrograph of our sample synaptosome preparation. The TEM illustrates intact pre-(PRE) and post-synaptic (POST) elements (arrows). Scale = 500nm. **(B)** Quality control western blots for whole brain (WB), cytosol (CYT), and synaptosome (SYN) fractions and associated quantification in control tissue. 5µg of protein per lane. **(C-G)** Quantification of western blots in **(A)**. Significance detected through One-Way ANOVA with Tukey’s post hoc test for multiple comparisons. **(H)** Representative western blot of WB, CYT, and SYN total protein levels of TrkB in Aldh1l1-TrkB.T1 cWT and Aldh1l1-TrkB.T1 cKO animals. 5µg of protein per lane. **(I-J)** Quantification of TrkB.FL and TrkB.T1 across various lysates of Aldh1l1-TrkB.T1 cWT versus Aldh1l1-TrkB.T1 cKO samples. Blots indicate that both TrkB.FL (I) and TrkB.T1 (J) are highly enriched in synaptosome preparations. However, conditional deletion of TrkB.T1 in astrocytes significantly decreases TrkB.T1 expression at the synapse **(J)**, indicating that this protein is enriched in perisynaptic astrocyte processes. (n = 5 animals).

Excitingly, when examining TrkB expression in Aldh1l1-TrkB.T1 animals, our results indicated that SYN lysates are enriched in TrkB.T1 expression (17.17-fold increase, p < 0.0001; **Figure 2H-J**) in Aldh1l1-TrkB.T1 cWT animals. However, we observed 89% depletion of TrkB.T1 in SYN lysates of Aldh1l1-TrkB.T1 cKO animals relative to Aldh1l1-TrkB.T1 cWT animals (**Figure 2J**), further confirming that astrocytes are the primary cell type expressing TrkB.T1. Importantly, no change in expression of presumably neuronal TrkB.FL was observed (**Figure 2I**). These findings highlight the robust presence of the TrkB.T1 isoform at the tri-partite synapse, particularly within PAPs, emphasizing its critical role in enabling astrocytes to respond to neuron-derived BDNF and facilitating astrocyte-synapse interactions.

### TrkB.T1 is implicated in the formation of glutamatergic synapses in motor and barrel cortex

Having established the localization of TrkB.T1 in PAPs, we next sought to elucidate how BDNF/TrkB.T1 signaling influences the development of synapses *in vivo*. We performed immunohistochemistry and confocal imaging of GFP-expressing astrocytes and probed for pre-synaptic vesicular glutamate transporters (VGluT1) and post-synaptic PSD-95 in layer II/III of the motor cortex (enriched in VGluT1-expressing corticocortical connections). We also employed a membrane-tagged Lck-GFP expressed under the control of the GfaABC1d promoter, allowing for the visualization of astrocyte morphology and neuropil infiltration of PAPs [1, 39]. Imaris analysis of astrocyte surface rendering and synaptic puncta detection allowed for a distance-based criterion for quantifying synapses (**Figure 3A**) [26]. Here, a 10 by 10 µm^3^ region of interest (ROI) containing astrocyte signal was isolated and surface rendering was applied to the astrocyte membrane-tethered eGFP signal. Next, the Spots Module in Imaris was employed to isolate VGluT1+ puncta and PSD+ puncta, allowing for a distance-based approach to quantify puncta and synapses – or colocalized pre- and post-synaptic elements, encapsulated within 1 µm of the astrocyte membrane (**Figure 3A**). We performed a mixed effects 2-way ANOVA to compare the effect of development and genotype on both the number of pre- and post-synaptic puncta and the number of colocalization events.

**Figure 3:**
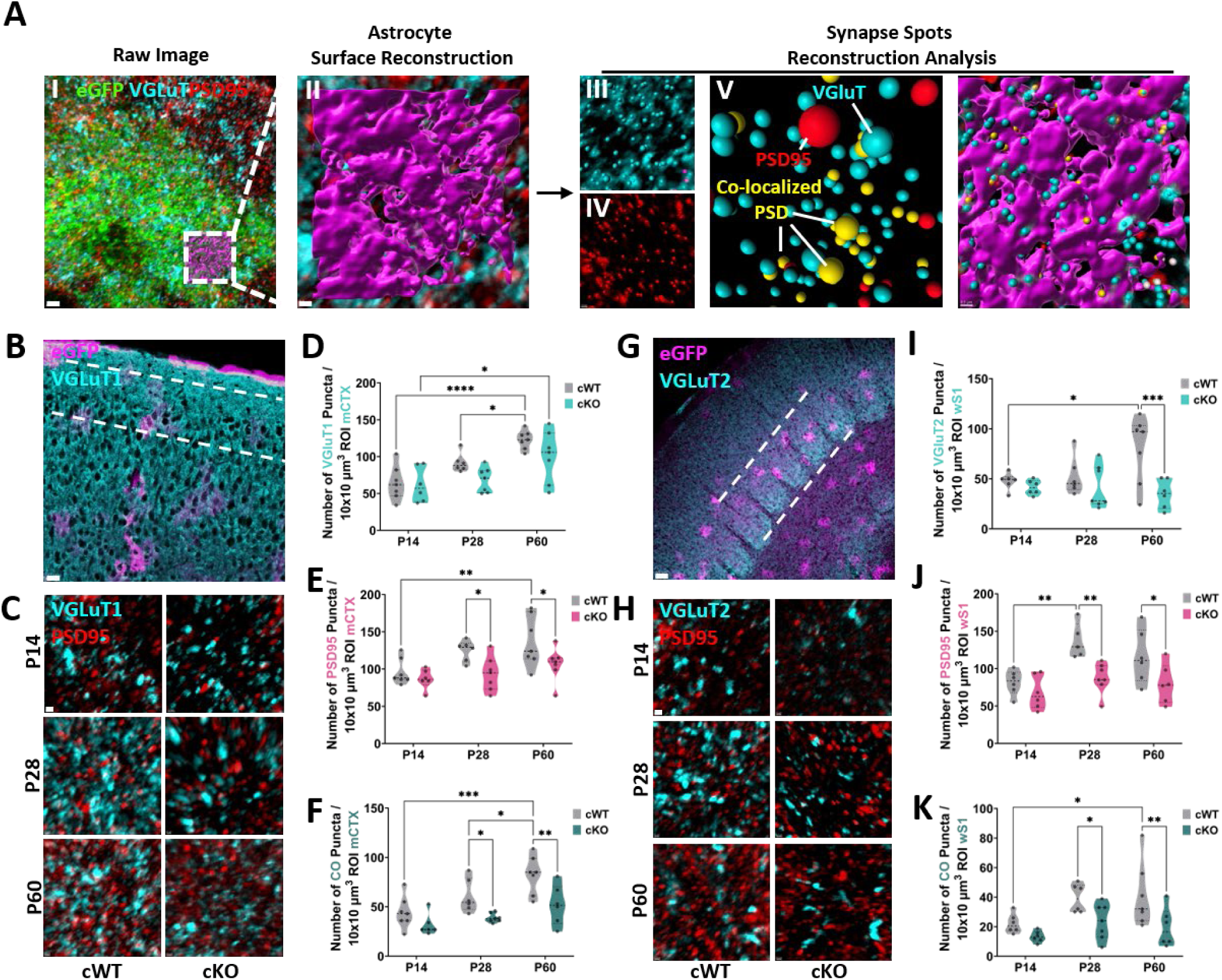
Conditional deletion of TrkB.T1 inhibits synapse formation in layer II/III of motor cortex and layer IV barrels of barrel cortex. **(A)** Representative images of Imaris workflow to analyze pre- and post-synaptic elements. *(I)* Confocal stack of example lck-eGFP expressing astrocyte and IHC for VGluT1 and PSD95. Dashed square indicates *(II)* selected region of interest (10×10μm3) within the astrocytic territory used to create a 3D rendering of the astrocyte’s surface. *(III)* Raw synaptic fluorescent signal of VGluT and PSD95 *(IV)* with 3D renderings of synaptic spots, depicted as cyan spheres for individual VGluT puncta and red spheres for individual PSD95 puncta. Puncta creation parameters ranged from 0.3 - 0.5μm estimated diameters. *(V)* Synaptic spots in the ROI. cyan dots are VGluT puncta, red spheres are PSD95 puncta, and yellow spheres indicate PSD95 puncta co-localized and within .75μm of a VGluT spot. High-throughput analysis allowed for the quantification of the total number of individual VGluT, PSD95, and co-localized puncta within ROIs. (I: Scale Bar = 2 μm; II-VI: Scale Bar = .5 μm). **(B)** Representative image of a coronal slice containing motor cortex. Dashed lines indicate region of interest. Scale Bar = 30 μm. **(C)** Representative IHC images of VGluT1 and PSD95 of Aldh1l1-TrkB.T1 cWT and Aldh1l1-TrkB.T1 cKO animals across development. Scale Bar = .5 μm. **(D)** VGluT1 expression increases between PND 14 and PND 60 for both Aldh1l1-TrkB.T1 cWT and Aldh1l1-TrkB.T1 cKO animals. **(E)** Deletion of TrkB.T1 results in a reduction of PSD95 puncta at PND 28 and PND 60. A significant increase in the number of puncta occurs between PND 14 and PND 60 in Aldh1l1-TrkB.T1 cWT, but not Aldh1l1-TrkB.T1 cKO mice. **(F)** The number of glutamatergic synapses, co-localized VGlutT1/PSD95, increases across development in Aldh1l1-TrkB.T1 cWT animals. In Aldh1l1-TrkB.T1 cKO animals, there is a delay and only a trending increase in synapse numbers. At PND 28 and PND 60, there is a significant reduction in synapse numbers in Aldh1l1-TrkB.T1 cKO animals. **(G)** Representative image of coronal slice containing barrel cortex. Dashed lines indicate region of interest. Scale Bar = 50 μm. **(H)** Representative IHC images of VGLuT2 and PSD95 of Aldh1l1-TrkB.T1 cWT and Aldh1l1-TrkB.T1 cKO animals across development. Scale Bar = .5 μm. **(I)** VGluT2 expression increases between PND 14 and PND 60 in Aldh1l1-TrkB.T1 cWT animals and remains steady in Aldh1l1-TrkB.T1 cKO animals. By PND 60 the number of VGluT2 puncta are significantly lower in Aldh1l1-TrkB.T1 cKO animals. **(J)** PSD95 in the barrel cortex significantly increases in Aldh1l1-TrkB.T1 cWT between PND 14 and PND 28 and stabilizes and lowers by PND 60. Its expression remains steady in Aldh1l1-TrkB.T1 cKO animals across development, resulting in significant reductions by PND 28 and PND 60 when compared to Aldh1l1-TrkB.T1 cWT groups. **(K)** The number of glutamatergic synapses in layer IV of the barrel cortex significantly increases across development in Aldh1l1-TrkB.T1 cWT animals but remains unchanged in the Aldh1l1-TrkB.T1 cKO animals. This results in significant reductions in the number of synapses in Aldh1l1-TrkB.T1 cKO animals when compared to Aldh1l1-TrkB.T1 cWT age-matched controls. Data are mean ± s.e.m. Mixed model ordinary two-way ANOVA with Turkey’s multiple comparisons test. Each dot represents an individual animal with n = 5-7 animals and n = 9-12 ROIs per animal (**D-F; I-K**). *p < 0.05, **p < 0.001, *** p < 0.001.

We imaged the motor cortex to examine if, at baseline, there are changes in synapse density in Aldh1l1-TrkB.T1 cKO mice. In layer II/III of the motor cortex (**Figure 3B**), we observed a significant effect based on genotype for VGluT1+ puncta (F(1, 34) = 5.143, p < 0.05; **Figure 3 B-D**), PSD95+ puncta (F(1, 34) = 11.20, p < 0.005; **Figure 3E**) and colocalized puncta (F(1, 33) = 18.48, p < 0.0005; **Figure 3F**). We also observed a significant effect on development for all three synaptic components (VGluT1: F(2, 34) = 16.54, p < 0.0001; PSD95: F(2, 34) = 6.291, p < 0.005; CO: F(2, 32) = 11.33, p < 0.0002). *Post hoc* Turkey’s multiple comparison tests found significant developmental increases in VGluT1+ puncta for both Aldh1l1-TrkB.T1 cWT (PND 14-PND 60: p < 0.0001, 95% C.I. = −87.69, −29.46; PND 28 – PND 60: p < 0.05, 95% C.I. = −1.576, 46.72) and to Aldh1l1-TrkB.T1 cKO animals (P14 - PND 60: p < 0.05, 95% C.I. = −68.59, −7.981; PND 28 - PND 60: p < 0.05, 95% C.I. = −60.83, −2.598). Importantly, the mean number of colocalized synaptic puncta was significantly higher in Aldh1l1-TrkB.T1 cWT animals at PND 28 (p < 0.05, 95% C.I. = 4.292, 39.66) and PND 60 (p < 0.005, 95% C.I. = 12.60, 47.97) when compared to Aldh1l1-TrkB.T1 cKO animals. As expected, the number of synapses in Aldh1l1-TrkB.T1 cWT animals significantly increased between PND 14 and PND 60 (p < 0.0005, 95% C.I. = −37.88, 4.781) and between PND 28 and PND 60 (p < 0.05, 95% C.I. = −42.78, −0.1233). In contrast, no significant increase in the number of colocalized puncta/synapses was observed across development in Aldh1l1-TrkB.T1 cKO animals.

To address if these changes were restricted to the motor cortex, we also evaluated the somatosensory cortex and focused on layer IV of the barrel cortex, enriched in VGluT2+ thalamacortical synapses. In the barrel cortex (**Figure 3G**), we observed a significant effect based on genotype for VGluT2+ puncta (F(1, 32) = 9.688, p < 0.005; **Figure 3G-I**) and PSD95+ puncta (F(1, 30) = 15.72, p < 0.0005; **Figure 3J**). We observed a significant effect in the number of synapses in the barrel cortex, across genotypes (F(1, 32) = 13.28, p < 0.0005; **Figure 3F**) and development (F(2, 32) = 4.274, p < 0.05; **Figure 3J**). *Post hoc* Turkey’s multiple comparison tests found that the mean number of VGluT2+ puncta was significantly higher in Aldh1l1-TrkB.T1 cWT relative to Aldh1l1-TrkB.T1 cKO animals at PND 60 (p = 0.0004, 95% C.I. = 21.74, 67.64), but not at other timepoints (**Figure 3I**). No significant upregulation of VGluT2 or PSD95 puncta was observed for Aldh1l1-TrkB.T1 cKO animals, but a significant increase in VGluT2 and PSD95 were observed for Aldh1l1-TrkB.T1 cWT animals between PND 14 (early postnatal), PND 28 (juvenile) and PND 60 (early adult) timepoints. This indicates that in the barrel cortex, deletion of TrkB.T1 disrupts the developmental increase of pre- and post-synaptic elements, resulting in a significant decrease in co-localized puncta in juvenile and early adult Aldh1l1-TrkB.T1 cKO animals (PND 28: p = 0.0351, 95% C.I. = 1.170, 30.35; PND 60: p = 0.0058, 95% C.I. = 6.599, 35.78). Altogether, our data suggests that during early developmental critical time periods when astrocytes typically infiltrate the neuropil [16], removal of TrkB.T1 in astrocytes results in dysregulated formation of pre- and post-synaptic elements in motor and barrel cortex. Consequently, this leads to a loss of synapses by PND 28 and 60.

### Astrocyte TrkB.T1 mediates astrocyte-synapse interactions across development

Our results indicate an important role for TrkB.T1 in the formation of synapses, yet, whether the lack of synapses in Aldh1l1-TrkB.T1 cKO animals is due to a change in astrocyte-synapse interactions is not known. As astrocytes mature morphologically, they extend leaflets to ensheathe synapses, forming PAPs. These PAPs exhibit structural plasticity influenced by experience-dependent plasticity [40–43]. While glutamate-induced activation of metabotropic glutamate receptors (mGluR1 and mGluR5) in PAPs triggers rapid structural changes [40], indicating glutamate is a driver triggering PAP plasticity, few studies have examined the molecular drivers of PAP-synapse interactions during development, with most focusing on experience-dependent plasticity or LTP models [43]. Our previous work has established BDNF/TrkB.T1 signaling in astrocyte morphological complexity [21]. Thus, we next assessed the developmental effects on synapse associations with astrocytes. Here, the *Surface Module* in Imaris was used to generate a 3D reconstruction from the astrocyte Lck-GFP signal and these reconstructions were used to create a masked channel isolating each astrocyte. Colocalization analysis was then performed between the masked Lck-GFP and VGluT1/2 or PSD95 voxels [29]. This served as an evaluation of astrocyte-synapse associations.

In layer II/III of the motor cortex, a mixed model 2way ANOVA revealed a significant effect based on development and genotype for VGluT1 (Development: F(2, 31) = 9.513, p < 0.005; Genotype: F(1, 31) = 49.36, p < 0.0005; **Figure 4A-C**) and PSD95 (development: F(2, 30) = 17.29, p < 0.0005; genotype: F(1, 30) = 33.55, p < 0.0001; **Figure 4D**). We also measured astrocytic volumes, as a measure of astrocyte morphological complexity and maturation. We observed a significant effect based on development (F(2, 31) = 8.548, p < 0.005) and genotype (F(1, 31) = 24.62 = 24.62, p < 0.0005) on astrocyte volumes as well as an interaction (F(2, 31) = 3.883, p < 0.05; **Figure 4E**). Across all developmental time points, multiple comparisons tests revealed a higher mean for the percentage of VGluT1 (PND 14: p < 0.005, 95% C.I. = 2.846, 14.94; PND 28: p < 0.0005, 95% C.I. = 8.069, 21.23; PND 60: p < 0.0005, 95% C.I. = 7.582, 19.68) and PSD95 (PND 14: p < 0.05, 95% C.I. = 0.93, 7.19; PND 28: p < 0.005, 95% C.I. = 1.99, 9.10; PND 60: p < 0.0005, 95% C.I. = 3.366, 9.626) associations in Aldh1l1-TRkB.T1 cWT astrocytes compared to Aldh1l1-cKO astrocytes. Overall astrocytic volumes were observed to significantly increase between PND 14 and PND 28 in Aldh1l1-TrkB.T1 cWT animals (p < 0.001), with no additional significant increases between PND 28 and PND 60 (p = 0.987) and an overall significant increase between PND 14 and PND 60 (p < 0.001).

**Figure 4:**
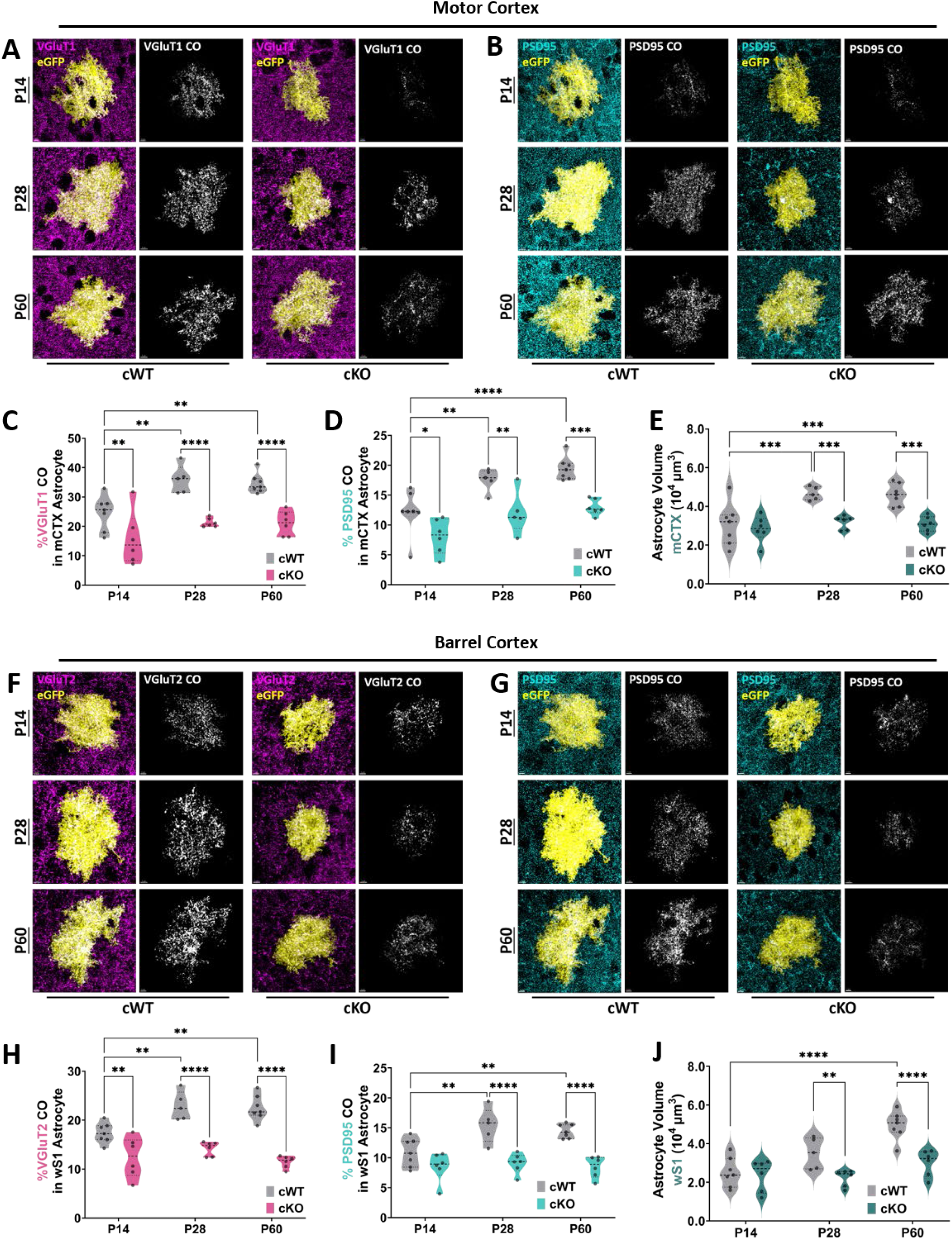
Conditional deletion of TrkB.T1 in astrocytes reduces astrocyte-synapse interactions. **(A)** Representative images of isolated Lck-GFP expressing astrocyte with pre-synaptic marker VGluT1 and **(B)** post-synaptic marker PSD95 in layer II/III motor cortex. When the two channels are merged, colocalized voxels (white) are quantified to determine the percent of astrocyte membrane colocalized with **(A)** VGluT1 and **(B)** PSD95. Images depict Aldh1l1-TrkB.T1 cWT and Aldh1l1-TrkB.T1 cKO raw Z-series images and their corresponding colocalized voxels across development. Scale Bar = 5 μm. **(C)** Astrocyte/VGluT1 contacts are significantly lower in Aldh1l1-TrkB.T1 cKO astrocytes across all stages in early development. The associations between PAP elements and VGluT1 increases in Aldh1l1-TrkB.T1 cWT animals but remain stable in Aldh1l1-TrkB.T1 cKO animals. **(D)** Astrocyte contacts with post-synaptic element PSD95 are significantly lower in Aldh1l1-TrkB.T1 cKO astrocytes across all stages in development. In Aldh1l1-TrkB.T1 cWT animals, astrocytes increase their associations with post-synaptic elements across developmental timepoints, but associations only trend towards an increase in Aldh1l1-TrkB.T1 cKO animals. **(E)** Overall morphometric properties of astrocytes increase across normal development, but are inhibited and lower in Aldh1l1-TrkB.T1 cKO animals. **(F)** Representative images of isolated Lck-GFP expressing astrocyte with pre-synaptic marker VGluT2 and **(G)** post-synaptic marker PSD95 in layer IV barrel cortex. When the two channels are merged, colocalized voxels (white) are quantified to determine the percent of astrocyte membrane colocalized with **(F)** VGluT2 and **(G)** PSD95. Images depict Aldh1l1-TrkB.T1 cWT and Aldh1l1-TrkB.T1 cKO raw Z-series images and their corresponding colocalized voxels across development. Scale Bar = 5 μm. **(H)** PAP and VGluT2 interactions significantly increase across development in normal conditions, but are consistently and significantly lower in Aldh1l1-TrkB.T1 cKO animals. **(I)** PAP and PSD95 interactions significantly increase across development in normal conditions, but are consistently and significantly lower in Aldh1l1-TrkB.T1 cKO animals. **(J)** Overall astrocyte volumes significantly increase between PND 14 and PND 60 in Aldh1l1-TrkB.T1 cWT animals, with no increase between PND 14 and PND 28. Morphometric properties remain constant in Aldh1l1-TrkB.T1 cKO animals and are significantly lower across all time points when compared to Aldh1l1-TrkB.T1 cWT controls. Data are mean ± s.e.m. Mixed model ordinary two-way ANOVA with Turkey’s multiple comparisons test. Each dot represents an individual animal with n = 5-7 animals and n = 3-4 astrocytes per animal (C-E; H-I). *p < 0.05, **p < 0.001, *** p < 0.001.

In the barrel cortex, a mixed model 2way ANOVA revealed a significant effect based on development and genotype for VGluT2 as well as an interaction (development: F(2, 31) = 6.179, p < 0.005; genotype: F(1, 31) = 94.29, p < 0.0005; Interaction: F(2, 31) = 4.573, p < 0.05; **Figure 4F & H**). We observed a similar result for PSD95 (development: F(2, 31) = 4.787, p < 0.05; genotype: F(2, 31) = 50.24, p < 0.0005; interaction: F(2, 31) = 3.507, p < 0.05; **Figure 4G & I**). We found a significant effect based on development and genotype and an interaction for astrocyte morphological maturation (Development: F(2, 31) = 16.11, p < 0.0005; Genotype: F(1, 31) = 22.75, p < 0.0005; Interaction: F(2, 31) = 5.222, p < 0.05; **Figure 4J**). Turkey’s multiple comparisons tests revealed significantly higher VGluT2 localization in Aldh1l1-TrkB.T1 cWT astrocytes across all developmental timepoints (PND 14: p < 0.005, 95% C.I. = 2.082, 7.893; PND 28: p < 0.0005, 95% C.I. = 5.499, 11.82; PND 60: p < 0.0001, 95% C.I. = 8.134, 13.95). Significantly higher levels of PSD95 and astrocyte associations in Aldh1l1-TrkB.T1 cWT animals were observed at later timepoints (PND 28: p < 0.0005, 95% C.I. = 3.7772, 8.931; PND 60: p < 0.0001, C.I. = 3.626, 8.366).

Between PND 28 and PND 60, as we had observed previously in global TrkB.T1^-/-^ animals [1], the mean astrocyte volume significantly increases in Aldh1l1-TrkB.T1 cWT animals (p < 0.005) and between PND 14 and PND 60 (p < 0.0001). Given that the barrel cortex develops earlier, with the critical period of layer IV synapses ending at PND14 [44], we were not surprised to find no significant increases in astrocyte volumes between PND14 and PND 60 in Aldh1l1-TrkB.T1 cWT animals. Importantly, we did not observe any significant increases in barrel cortex astrocyte volumes in Aldh1l1-TrkB.T1 cKO animals Our data suggest that BDNF and TrkB.T1 signaling in development is an important signaling mechanism that mediates astrocyte-synapse interactions and morphological maturation. To address if these synaptic abnormalities were a result of reduced astrocyte engagement, we quantified the mRNA expression of common astrocyte-derived synaptogenic cues in the somatosensory cortex of Aldh1l1-TrkB.T1 cWT and Aldh1l1-TrkB.T1 cKO animals. Interestingly, we found reduced expression of the synapse promoting cues *Thbs1* and *Sparcl1*, and increased expression of the astrocyte-derived synaptic pruning factor *Megf10* at different timepoints in development (**Supplemental Figure 2**). We also observed significant reductions in PAP-related genes such as *Slc1a3*, Slc1a2, and *Kcnj10* (**Supplemental Figure 2H**). These results indicate that the loss of synapses may be due to reduced astrocyte interactions at the synapse, decreasing the ability for astrocytes to release or express synapse-promoting genes, or genes related to their ability to maintain ion and neurotransmitter homeostasis.

Collectively, our findings from *in vitro* co-cultures, synaptosome preparations, and *in vivo* three dimensional confocal microscopy converge to establish TrkB.T1 as a critical mediator of astrocyte-synapse interactions, demonstrating an enrichment in PAPs and its essential role in maintaining glutamatergic synapses and astrocyte-synapse interactions in cortex, thereby underscoring the fundamental importance of astrocytic BDNF/TrkB.T1 signaling in shaping synaptic landscapes in the mammalian brain.

## DISCUSSION

In the present study, we uncover a pivotal role for astrocytic TrkB.T1 in orchestrating the development of excitatory synapses. Through neuron-astrocyte co-cultures, we demonstrated that TrkB.T1 KO astrocytes show a marked reduction in PAPs and weakened interactions with excitatory synapses. Furthermore, the astrocyte-specific deletion of TrkB.T1 *in vivo* led to significant deficits in excitatory synapse numbers and astrocyte-synapse associations in the motor and barrel cortex. These impairments became evident during the later stages of postnatal development, aligning with critical periods of synaptic pruning and maturation.

Proper synapse development and maintenance are critical for normal brain function and regulation of behavior. BDNF/TrkB.FL signaling in neurons is considered a key mechanism underlying synapse formation, maturation, and plasticity by acting on both pre- and post-synaptic elements [45]. For example, BDNF/TrkB signaling enhances quantal neurotransmitter release [46], regulates the recruitment of synaptic scaffolding proteins like PSD95 at postsynaptic elements [47], modulates both AMPA and NMDA receptor trafficking and function [48–50], and stimulates structural dendritic spine dynamics [51–53]. Due to its canonical role at the synapse, it is not surprising that disruptions in this signaling pathway have been implicated in numerous neurodevelopmental and neuropsychiatric disorders [54, 55]. However, unlike TrkB.FL, the role and mechanisms of TrkB.T1 at the synapse has been less studied. Given the fact that TrkB.T1 is the predominant isoform expressed in both mouse [2, 4] and human cortex [5, 6], elucidating the downstream mechanisms is of critical importance for the understanding of BDNF signaling at the synapse.

In addition to BDNF/TrkB signaling, astrocytes are another major contributor to the development and maturation of synapses. Astrocytes release synaptogenic cues including hevin and sparc [56], thrombospondins [57], glypicans 4/6 [58], and TGF-β [59, 60] to induce the formation of synapses and stabilize new ones. Astrocyte cell adhesion molecules are an additional mechanism through which astrocytes promote synapse formation. For example, astrocyte ephrin-A3 (EphA3) [61] physically contacts postsynaptic elements by interacting with neuronal EphA4, and this engagement matures dendritic spine morphology [62]. Astrocytic Necl2 with axonal Necl3 [63] as well as Connexin30 in astrocytes [64] are additional contact-mediated mechanisms that have been implicated in both astrocyte morphogenesis and synapse formation. In addition, homophilic interactions between astrocyte γ-protocadherins (γ-Pcdh) and neuronal γ-Pcdh isoforms are critical for synapse formation and stabilization during early development [65]. Collectively, these studies and many others demonstrate the importance for physical interactions between neurons and astrocytes at the synapse for the proper development of synapses and for the induction of astrocyte morphological complexity. However, there is still very limited knowledge regarding how and when PAPs arrive at the synapse. Our work has previously demonstrated that astrocytes predominantly and nearly exclusively express TrkB.T1 and rely on BDNF/TrkB.T1 signaling to undergo their morphological maturation [1]. Furthermore, we demonstrated that *in vitro*, deletion of TrkB.T1 resulted in aberrant synapse formation and function.

To begin to dissect cellular mechanisms underlying the loss of synapses following deletion of TrkB.T1, we used neuron-astrocyte co-cultures to examine the temporal and spatial formation of PAPs around excitatory synapses. We confirmed previous results demonstrating WT astrocytes enhance the development of synapses *in vitro* [1, 57, 66] and revealed neuronal morphological complexity enhanced astrocyte morphological features. This finding confirms studies that have identified neuron-astrocyte communications as important mediators underlying astrocyte morphogenesis [39, 67, 68]. We visualized astrocyte morphology *in vitro* utilizing GLAST to reveal the astrocyte membrane and more closely examine astrocyte-synapse associations. Relative to the cytoskeletal-associated GFAP, GLAST reveals up to four-fold more astrocyte membrane [69] and PAPs have been shown to highly express GLAST [70]. Due to this, we considered the high-intensity GLAST immunofluorescence and circular structures observed surrounding colocalized pre- and post-synaptic puncta to be PAP-like structures. The marked reduction in both synapse numbers and PAPs in TrkB.T1^-/-^ KO conditions suggest a crucial role of TrkB.T1 in mediating astrocyte-synapse interactions.

Given that BDNF is necessary for the development of synapses and that it can be expressed following exposure to KCl *in vitro* [36, 71], our findings indicate that the highly expressed TrkB.T1 in PAPs allows astrocytes to respond to local BDNF. At an active and maturing synapse, BDNF is released locally for structural and functional synapse formation [72], our data suggests that BDNF signaling to astrocyte TrkB.T1 may trigger the attraction of astrocyte leaflets to these nascent synapses, where they can engage with synaptic elements, become PAPs, and further mature the structural and functional properties of the synapse.

*In vivo*, conditional deletion of TrkB.T1 in astrocytes at the onset of synaptogenesis resulted in significant reductions in the development of glutamatergic synapses in the motor and barrel cortex. Furthermore, we demonstrated that TrkB.T1 deletion in astrocytes reduced astrocyte-synapse interactions. We observed significant reductions in synapse numbers and astrocyte-synapse interactions beginning at PND 28, coinciding with peak BDNF expression [1, 34]. Interestingly, in the ventromedial hypothalamus, BDNF/TrkB.T1 signaling regulates PAP invasion into synapses, with BDNF depletion increasing astrocytic process invasion and GLT-1 expression [73]. This work suggests that *in vivo,* BDNF/TrkB.T1 does serve to mediate astrocyte-synapse interactions in regions such as the VTA, which undergoes plasticity in response to levels of BDNF driven by satiety [73]. Furthermore, in a mouse model of temporal lobe epilepsy, modulation of astrocytic BDNF and TrkB have been shown to significantly affect neuronal activity and cognitive function [74], indicating an overall important of astrocyte BDNF/TrkB in overall activity and behavior. Our *in vitro* work suggests that during both development and high neuronal activity, BDNF is indeed a trigger of astrocyte plasticity through TrkB.T1. Indeed, early work in the field demonstrated that TrkB.T1 mediates Ca^2+^ signaling in astrocytes in response to BDNF [75]. Furthermore, TrkB.T1 in cultured astrocytes has been shown to mediate astrocyte plasticity via Rho GDP dissociation Inhibitor 1, presumably trough actin dynamics [76] that may be activated through Ca^2+^ signaling. Although our current work is limited in the temporal resolution required to observe immediate structural plasticity, it offers insights into the role of TrkB.T1 in allowing astrocytes to engage with maturing synapses across time.

In the context of the cortex, our data indicates that loss of TrkB.T1 in astrocytes may render them unable to respond effectively to the developmental increase of BDNF expression, consequently inhibiting their interactions at developing synapses. This impairment likely affects the expression of synapse-promoting cues, ion channels, and neurotransmitters critical for astrocytic support of synapses [56, 57, 60, 65, 68]. We observed decreases in mRNA expression of GLT-1, GLAST, and Kir4.1, presumably due to reduced PAP formation, which may contribute to the reduction in synapse numbers observed. In addition, it is important to note that glutamate-induced activation of metabotropic receptors (mGluR1 and mGluR5) in PAPs triggers rapid structural changes in PAPs in the context of experience-dependent plasticity [40, 43], suggesting glutamate is an important driver of PAP plasticity. Herein, we propose that BDNF signaling and TrkB.T1 work akin to glutamate signaling in PAPs, allowing them to engage in the structural organization required for them to engage at the synapse, where they can express the necessary synapse-promoting and synapse homeostatic genes and proteins required to properly mature a synapse. With that said, a more comprehensive proteomic analysis could provide deeper insights into the molecular mechanisms underlying these predictions. Overall, our results suggest that TrkB.T1 is a prerequisite protein for astrocytes to engage with synapses, thus allowing for strong support, synapse maturation, and potentially facilitating synaptic plasticity.

It is important to acknowledge the limitations of our study, notably the use of confocal microscopy rather than higher resolution techniques like electron microscopy. PAPs are nanoscale structures (10-100 nm in diameter) that are difficult to visualize using conventional light microscopy due to diffraction limits [77]. Volume electron microscopy has been a very valuable tool to more closely examine the associations between PAPs and synapses [78]. However, confocal microscopy has been a valuable tool to obtain metrics such as astrocyte territory size [79], neuropil infiltration volumes [7, 68], and territory volumes [80, 81]. Further, the employment of viruses to express the membrane-tethered Lck domain fused to GFP has been a valuable approach to investigate astrocyte major branching organization and to form 3D reconstructions [1, 82]. The Lck-GFP reporter and its association with the astrocyte membrane enables uniform distribution across astrocyte membranes, providing trusted representation and reconstruction of astrocytes. In regards to astrocyte-synapse interactions, three-dimensional reconstructions of Lck-GFP astrocytes and the colocalization of this signal with synaptic elements has previously been used to study astrocyte-synapse interactions in the context of development [83], drug addiction [84–86], and stress [87], indicating that this is a conventional approach to study astrocyte-synapse associations and to capture changes that result from development or plasticity.

In conclusion, this study establishes a critical role for astrocytic TrkB.T1 signaling in the regulation of synapses and astrocyte-synapse associations during developmental time periods. Conditional deletion of TrkB.T1 in astrocytes reduced somatosensory and motor cortex synapse densities and astrocyte-synapse associations. Investigations using neuron-astrocyte co-cultures revealed impairments in the formation of PAPs and their interactions with excitatory synapses, providing insight into how this pathway governs astrocyte-mediated synaptic support and plasticity. These findings have implications for neurodevelopmental and neuropsychiatric disorders implicated by astrocytes and synapses. Ultimately, elucidating the role of BDNF and astrocyte TrkB.T1 signaling may unveil novel therapeutic avenues to target synaptic and behavioral abnormalities.

## ACKNOWLEDGEMENTS

This research project was funded by NIH/NINDS R01 NS120746 to MLO and NIH/NINDS F31 NS127511 to BTCP.

---

**Supplemental Figure 1:**
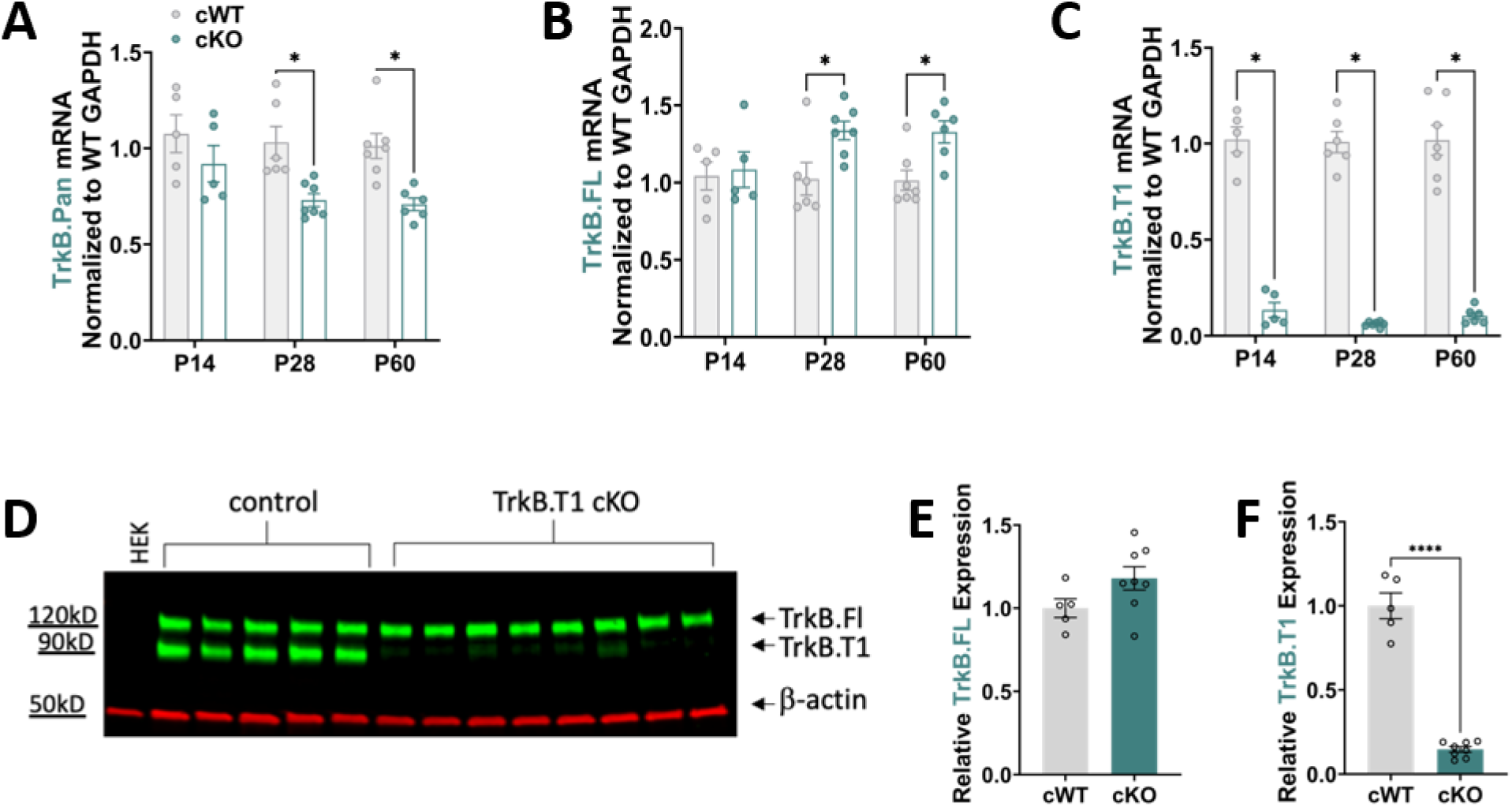
Effective knockdown of TrkB.T1 in Aldh1l1-TrkB.T1 cKO animals. **(A-C)** QPCR analysis of Aldh1l1-TrkB.T1 cWT and Aldh1l1-TrkB.T1 cKO somatosensory cortex samples. Relative to Aldh1l1-TrkB.T1 cWT samples, data demonstrates (A) decreased mRNA expression of TrkB at PND 28 and PND 60 **(B)** an increase in TrkB.FL expression at PND 28 (t(11) = 2.678, p < .05) and PND 60 (t(11) = 3.249, p < .05), and **(C)** significantly low expression of TrkB.T1 across all timepoints in Aldh1l1-TrkB.T1 cKO samples, indicating both effective and isoform specific tamoxifen-inducible knockdown across relevant timepoints. (n = 5-7 animals) **(D)** Representative western blot of total protein levels in whole cortex of TrkB in Aldh1l1-TrkB.T1 cWT and Aldh1l1-TrkB.T1 cKO animals. 5µg of protein per lane. **(E-F)** Quantification of TrkB.FL and TrkB.T1 in Aldh1l1-TrkB.T1 cWT versus Aldh1l1-TrkB.T1 cKO samples. (n = 5 cWT; n = 8 cKO).

**Supplemental Figure 2:**
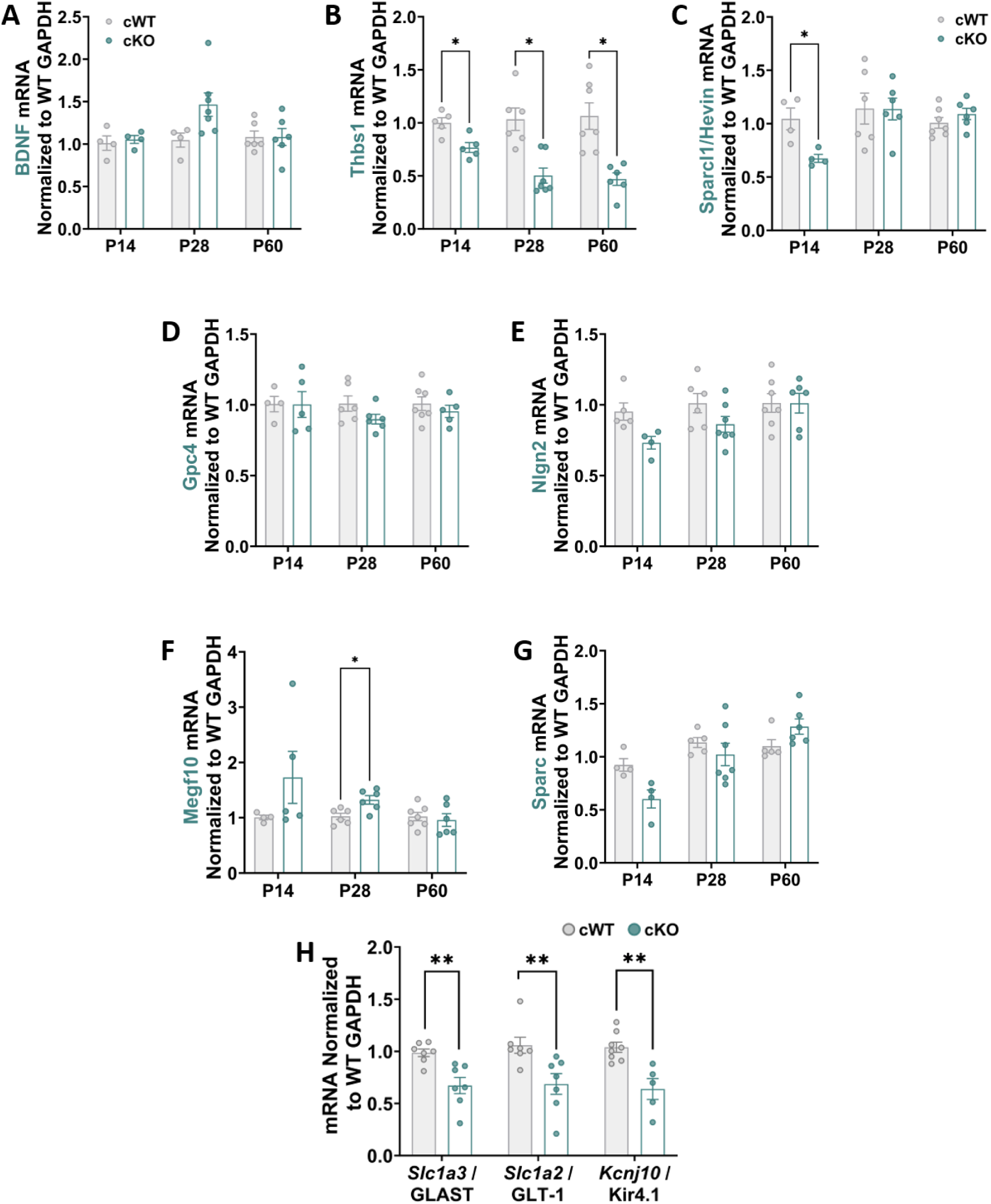
Synapse promoting and Synapse inhibiting cues across development. **(A-G)** qPCR analysis of Aldh1l1-TrkB.T1 cWT and Aldh1l1-TrkB.T1 cKO somatosensory cortex samples across development. **(B)** There was a significant reduction in *Thbs1* expression in Aldh1l1-TrkB.T1 cKO at all three time points. *Thbs1* codes for Thrombospondin, a synaptogenic cue released by astrocytes. **(C)** *Sparcl1/*Hevin was significantly downregulated at P14 I Aldh1l1-TrkB.T1 cKO animals, but not at any other timepoints. We observed no changes in **(A)** *Bdnf*, **(D)** *Gpc4,* or **(E)** *Nlg2* expression. Interestingly, **(F)** *Megf10* which is associated as an astrocyte-derived cue for synaptic pruning was higher in Aldh1l1-TrkB.T1 cKO mice relative to cWT. **(G)** No changes were observed in the expression of *Sparc*. **(H)** qPCR analysis of Aldh1l1-TrkB.T1 cWT and Aldh1l1-TrkB.T1 cKO somatosensory cortex samples at P28. We observed significant reductions in the expression of GLAST, GLT-1, and Kir4.1 mRNA.

